# Agonist-induced Functional Analysis and Cell Sorting, a novel tool to select and analyze neurons: Fragile X as a proof of concept

**DOI:** 10.1101/2020.04.20.050997

**Authors:** Sara Castagnola, Julie Cazareth, Kevin Lebrigand, Marielle Jarjat, Virginie Magnone, Sebastien Delhaye, Frederic Brau, Barbara Bardoni, Thomas Maurin

## Abstract

To get a better insight into the dynamic interaction between cells and their environment, we developed the agonist-induced Functional Analysis and Cell Sorting (aiFACS) technique, which allows the simultaneous recording and sorting of cells in real-time according to their immediate and individual response to a stimulus. By modulating the aiFACS selection parameters, testing different developmental times, using various stimuli and multiplying the analysis of readouts, it is possible to analyze cell populations of any tissue, including tumors. The association of aiFACS to single-cell transcriptomic allows to build a tissue cartography based on specific functional response/s of cells.

As proof of concept, we used aiFACS on the dissociated mouse brain, a highly heterogenous tissue, enriching it in interneurons upon stimulation with an agonist of the glutamate receptors and upon sorting based on calcium levels. Further single-cell RNA-seq of these aiFACS-selected interneurons resulted in a nine-cluster classification. Furthermore, we used aiFACS on interneurons derived from the brain of the *Fmr1*-KO mouse, a rodent model of Fragile X syndrome. We show here that these interneurons manifest a generalized defective pharmacological response compared to wild type, affecting all the analyzed cell clusters at one specific post-natal developmental time.

## INTRODUCTION

The selection, cloning and morphological/functional characterization of individual cells in a suspension or from a tissue is a fastidious and lengthy procedure, although several techniques have been set up to reach this goal: limiting dilutions, laser micro-dissection (Datta et al. 2015), cytoplasm aspiration (Cadwell et al. 2016; Fuzik et al. 2016) micro fluidics (Pollen et al. 2014; Tasic et al. 2016; Zeisel et al. 2015) and flow cytometry (Fulwyler 1965). The latter is a robust and powerful technique that allows the fast analysis and sorting of a subset of cells from an initial heterogeneous sample. Cells can be sorted according to various parameters and are amenable to further characterization such as single-cell transcriptomics or mass spectrometry. Single-cell transcriptomics is the unique tool that allows a precise and simultaneous analysis of the expression levels of thousands of genes in a complex heterogeneous population at the single-cell level (Poulin et al. 2016). Applied to the study of a pharmacological stimulation, this technique can decipher homeostatic gene regulation at the tissue level from the modulation of the abundance of a given cell population, a phenomenon that is widely observed during development (Ofengeim et al. 2017; Poulin et al. 2016).

Coupling flow cytometry analysis and cell sorting with single-cell transcriptomics makes it possible to link a cell phenotype to its genotype, providing a direct access to the molecular cues of a given phenotype. In this context, the key point is to submit the cells under analysis to the same stimulus. The functional FACS analyses commonly used to measure pH or calcium (Chow et al. 2001; Vines et al. 2010), for instance, consist in adding the drug into the sample tube containing the cell suspension and then analyze and sort them. It takes up to 30 seconds for the sample to reach the flow cell (the chamber in which the cells are aligned to pass one-by-one through the light beam for sensing), so we can consider that the system measures pH or calcium from probe fluorescence “*at equilibrium*”, since the measurement and the stimulus are not simultaneous. Indeed, commercial FACS or flow cytometry analyzers are not yet adapted to the analysis of differentiated individual calcium responses as observed in the field of view of a microscope, with variations of the fluorescence (of the calcium probes) occurring over intervals of a few milliseconds to seconds. In many commercial systems the cells in suspension in the sample tube are injected under pressure into the cytometer fluidics, which prevents the addition of any drug at atmospheric pressure requiring the removal of the sample tube for this operation. This, in turn, constrains the observation of “stationary” cellular responses rather than dynamic. Moreover, this method of stimulation presupposes that the first and last cells analyzed in the tube behave in an equivalent way. Although a technical approach that allows the addition of drugs into the sample tube without discontinuity has been proposed (Arnoldini et al. 2013; Vines et al. 2010), its realization requires all cells to be simultaneously exposed to the drug but necessarily analyzed at different times. Encouraging approaches to “rapid” calcium response measurements by flow cytometry have been proposed in the past, based on a fluidic change in a FACS analyzer (Tárnok 1996; Zwartz et al. 2011). A way to monitor brief kinetics and the possibility to sort the cells by modifying the tubing configuration of a FACStar Plus cytometer was described (Dunne 1991). However, no sorting evidence was demonstrated. These approaches remained in the form of prototypes and were only used to carry out analysis (Time zero Instrument). On a simpler and similar basis, we have performed an instrumental development on a cell sorter.

The agonist-induced Functional Analysis and Cell Sorting (aiFACS) prototype is developed on a FACS Aria III as an add-on. It allows to analyze all the aforementioned parameters but also to sort cells according to their immediate and individual response to a stimulus. We used it on mouse brains and we were able to obtain preparations enriched in interneurons. Further application of aiFACS to FXS mouse brains highlights the functional and molecular impairment of interneurons, thus validating in this model the power of this technique as a new tool to study brain disorders. Beside the brain, aiFACS can be employed to analyze cell populations of any normal and pathological tissue, including tumors.

## RESULTS

### Set up of Agonist-Induced Functional Analysis and Cell Sorting (aiFACS)

We designed the aiFACS prototype in order to be able to sort cells according to their response to a pharmacological stimulation that is monitored in real time with a fluorescent calcium indicator (Figure 1A). The sorter fluidics (BD FACAria III) is modified to allow drug injection into the sample line. Thanks to a “Y” connector, a tubing (shown in red, Figure 1A) with a diameter equivalent to that of the sample line (shown in blue, Figure 1A) is connected to two syringes, upstream of the solenoid valve and the flow cell. Two valves open or closed alternately control the injection syringes containing PBS and agonist and allow the injection pad to be selected. A peristaltic micro-pump makes it possible to control the speed of injection of the solutions in order to synchronize it with the FACS flowrate. The incubation time between each cell and the drug is thus controlled (red tubing, Figure 1A). In basal conditions, a sample resuspension buffer (D-PBS) is infused. To promote stimulation, the experimental drug, concentrated twice, is injected (2X agonist). The cells are sorted according to their response to the drug.

**FIGURE 1.**
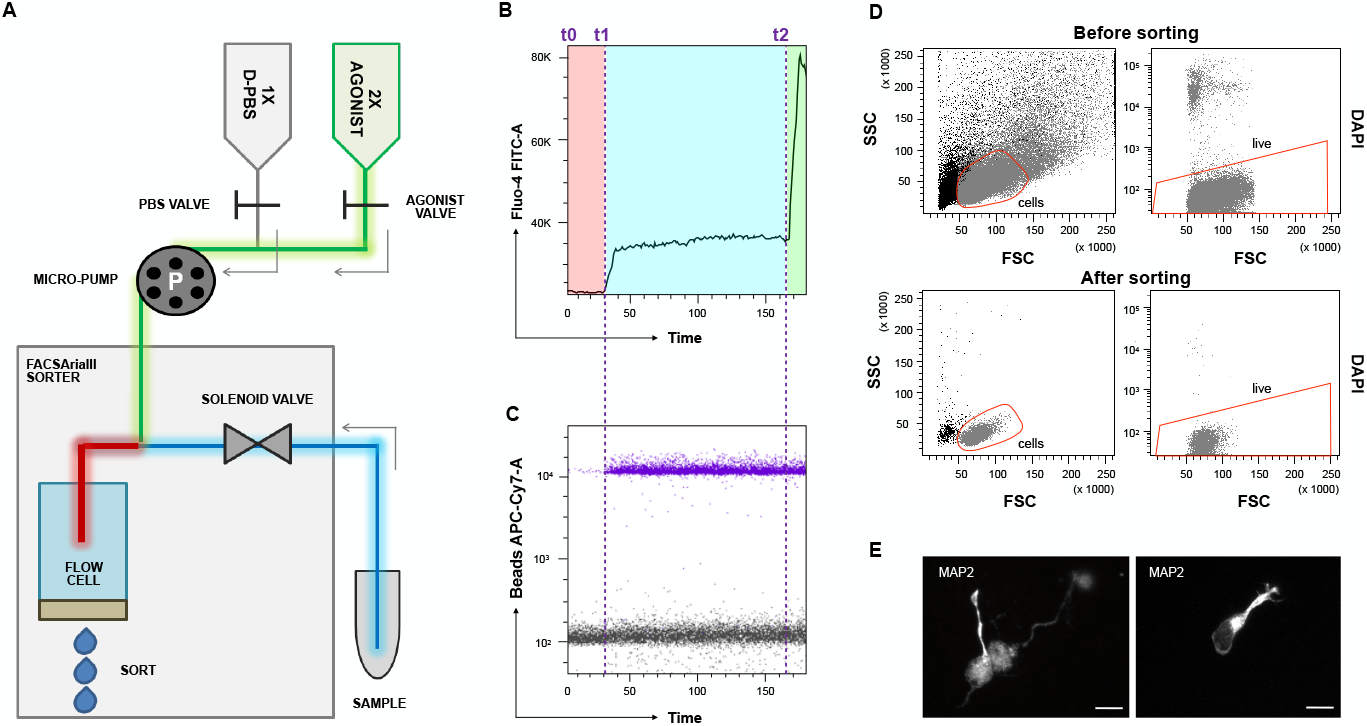
The aiFACS technique. **A)** Schema of the instrumental apparatus: BD FACSAria III implemented with the aiFACS device. The fluidics of the sorter is modified to allow the injection of a pharmacological agonist. Two syringes, one containing D-PBS (in grey) and the other containing the agonist concentrated twice (in green) are connected to their respective tubing: the D-PBS tubing (in grey) and the agonist tubing (in green). These are further connected to a downstream Y-shaped connector that enters the flow cell (the chamber in which the cells are aligned to pass one-by-one through the light beam for sensing). The sample is connected to the sorter through a tubing (in blue) having the same diameter as the agonist tubing (in green). A peristaltic micro-pump (P) allows to control the speed of solution injection and to synchronize it to the speed of sample flow in the sorter. The incubation time between each cell of the sample and the agonist is also controlled (red tubing). **B)** Time *vs* Fluo-4 AM bi-parametric graph showing the response of the cells to different stimuli in real time. At time t0 the opening of the D-PBS valve starts the perfusion and the baseline levels of fluorescence (in the red rectangle) are obtained with continuous perfusion. At time t1 the D-PBS valve is closed and that of the agonist is opened. The magnitude of the cellular calcium response to the KCl agonist (50 mM final) is shown in the light-blue rectangle. At time t2 ionomycin is added to KCl as a positive control of stimulation. The maximal response of the cells is displayed in the green rectangle. **C)** Addition of beads to the agonist solution allows real time detection of the agonist presence. Bead fluorescence is shown in purple. **D)** aiFACS allows viable recovery of stimulated cells. Upper panels: discrimination of cells based on scatter parameters (FSC: forward scatter; SSC: side scatter) before sorting (55.3 % of the total population; left panel). The pre-sort viability is determined by labelling the cells with DAPI (95.5 % of the cells in the red region; right panel). Lower panels: discrimination of cells based on scatter parameters (FSC: forward scatter; SSC: side scatter) after sorting (90.9 % of the total population; left panel). The viability of the cells after KCl stimulation and aiFACS sorting is determined by re-analyzing the DAPI staining (99,7 % of the cells in the red region; right panel). E) Sorted neurons are viable and can grow neurites when plated on L-ornitin-coated glass coverslips, cultivated for up to 6 Days *In Vitro* (DIV 6) in complete neurobasal medium and analyzed by immunocytochemistry (MAP2 staining). Scale bars: 10 μm.

Using the Miltenyi Neuron Isolation kit (Berl et al. 2017; Holt and Olsen 2016), we freshly dissociated 18 Post-Natal Day (PND 18) mouse adult brains to obtain RNA from the neuronal and non-neuronal collected populations that we used to measure the expression of several neuronal markers by RT-qPCR (Supplemental Figure S1). The neuron fraction is enriched in interneuron (*Gad2*, *Cnr1* not shown) and pyramidal cell (*Itpr1*) markers, and depleted in microglial (*Grn*), immature neuron (*Sox2*) and astrocyte (*Gfap*) markers. Oligodendrocytes (*Mog2*) and endothelial cells (*Rapgef4*) were not significantly removed using this magnetic separation procedure and they were not taken into account in the subsequent analyses.

We injected neurons labelled with Fluo-4 AM in order to monitor the calcium response and we set up the machine using the parameters indicated in red in Supplemental Figure S2A. In this first step, based on our previous experience with ratiometric calcium imaging (Castagnola et al. 2018), we used KCl (50 mM final) as agonistic drug to elicit calcium entry into cells. We added fluorescent beads to monitor the agonist perfusion and the increase in K^+^ ion concentration in the sample line. At time t0 the D-PBS perfusion starts and it is stopped at time t1, when the KCl perfusion is opened (Figure 1B). Each cell will be in contact with KCl for 3.2 seconds (Supplemental Figure S2A). The appearance of the beads corresponds to the arrival of cells stimulated with KCl in the flow cell (Figure 1C). In order to verify the amplitude of neuronal response and have a positive control of stimulation, ionomycin is added to KCl at time t2 (Figure 1B). Pre-sort viability is determined by labeling dissociated neurons with DAPI staining (Figure 1D). The viability of the cells having responded to KCl and after sorting by aiFACS can be monitored with the previous labeling and it is determined through re-analysis of the cells (Figure 1D). Remarkably, post-aiFACS KCl-responsive cells are viable (>90%) and capable of growing neurites when seeded on ornithine-coated glass coverslips (Figure 1E).

### aiFACS selection and characterization of AMPA-responding interneuron populations

To obtain a proof of concept that this new method allows to select and study neuronal populations using a pharmacological stimulation, we decided to activate a ionotropic receptor by its agonist and we used α-amino-3-hydroxy-5-methyl-4-isoxazolepropionic acid (AMPA). After brain dissociation, cells stained with Fluo-4 were gated based on their fluorescence. We injected the cells in the FACS supplying the flow cell with a neutral D-PBS (baseline condition). After 3 minutes of recording, we stimulated cells with an AMPA solution (100 μM final). Also in this case each cell was in contact with the drug for 3.2 seconds. In our analysis we focused on the cells which exhibit a homogenous Fluo-4 staining and we called them gating-dependent cells population 1 (GD1 cells; Figure 2A). This cell population is enriched in *Gad2*- and *Calb2*-expressing cells compared to the remaining cell population (GD2 cells; Figure 2B). This suggests that our strategy favored the selection of interneurons (Le Magueresse and Monyer 2013). To get more insight in the interneuron sub-population selected, GD1 cells were used to carry out single-cell transcriptomics with the 10x Genomics Chromium and Illumina sequencing platform. After quality control, we analyzed 2,170 cells (1,287 AMPA and 883 baseline), with a median UMI per cell of 6,652. The canonical correspondence analysis (CCA) of the two aggregated samples led to the identification of nine cell clusters (Figure 3A and 3B). The AMPA response of each cluster is illustrated in Figure 3C. GD1 cells are broadly split into inhibitory (*Dcx*, *Gad1*, *Gad2*, *Dlx6*, *Dlx1*, *Dlx5*, and *Dlx2*) and excitatory neurons (*Slc17a7*, *Slc17a6*, *Nrn1*, *Neurod1*, and *Neurod2*) (Figure 3A and 3B).

**FIGURE 2.**
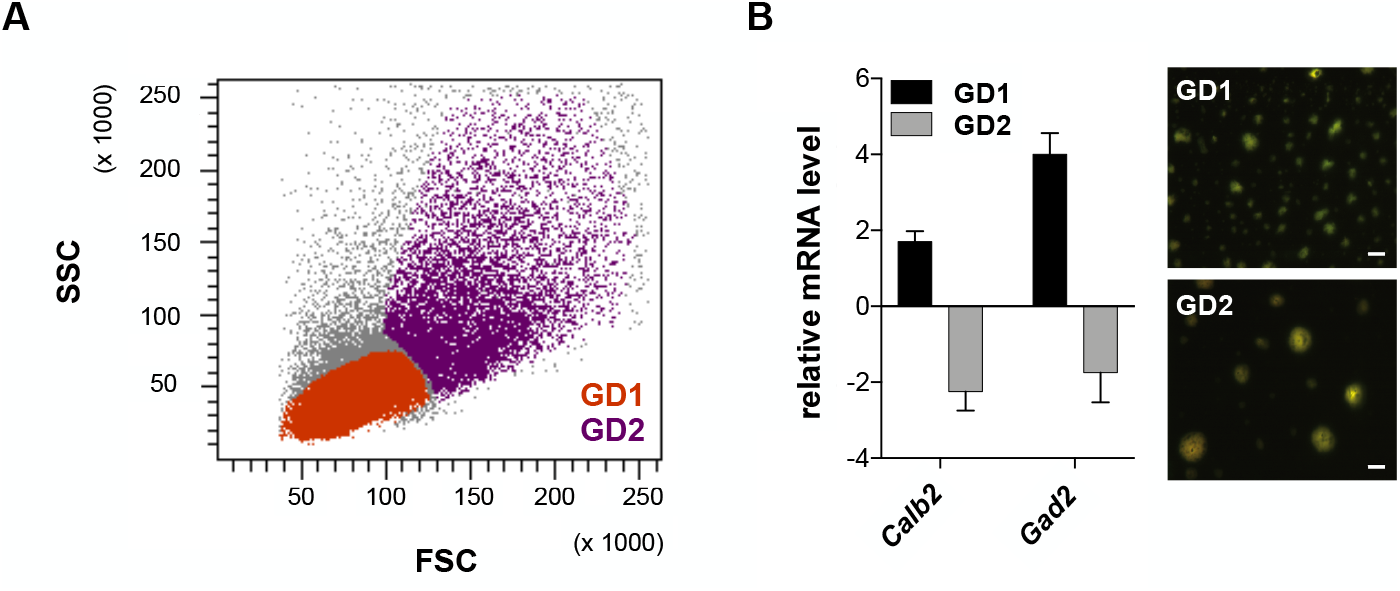
The gating strategy. **A)** Discrimination of cells based on scatter parameters (FSC: forward scatter; SSC: side scatter). Cells were gated according to their size/structure. GD1: gating-dependent cell population 1; GD2: gating-dependent cell population 2. **B)** RNA was purified from WT GD1 and GD2 cells. The inhibitory neuron marker levels were analyzed in both populations by RT-qPCR. The graph (on the left) shows mRNA expression relative to the total WT neuron suspension (WT input). GD1 and GD2 cells were live-imaged 2 hours after aiFACS sorting (on the right). Scale bars: 15 μm.

**FIGURE 3.**
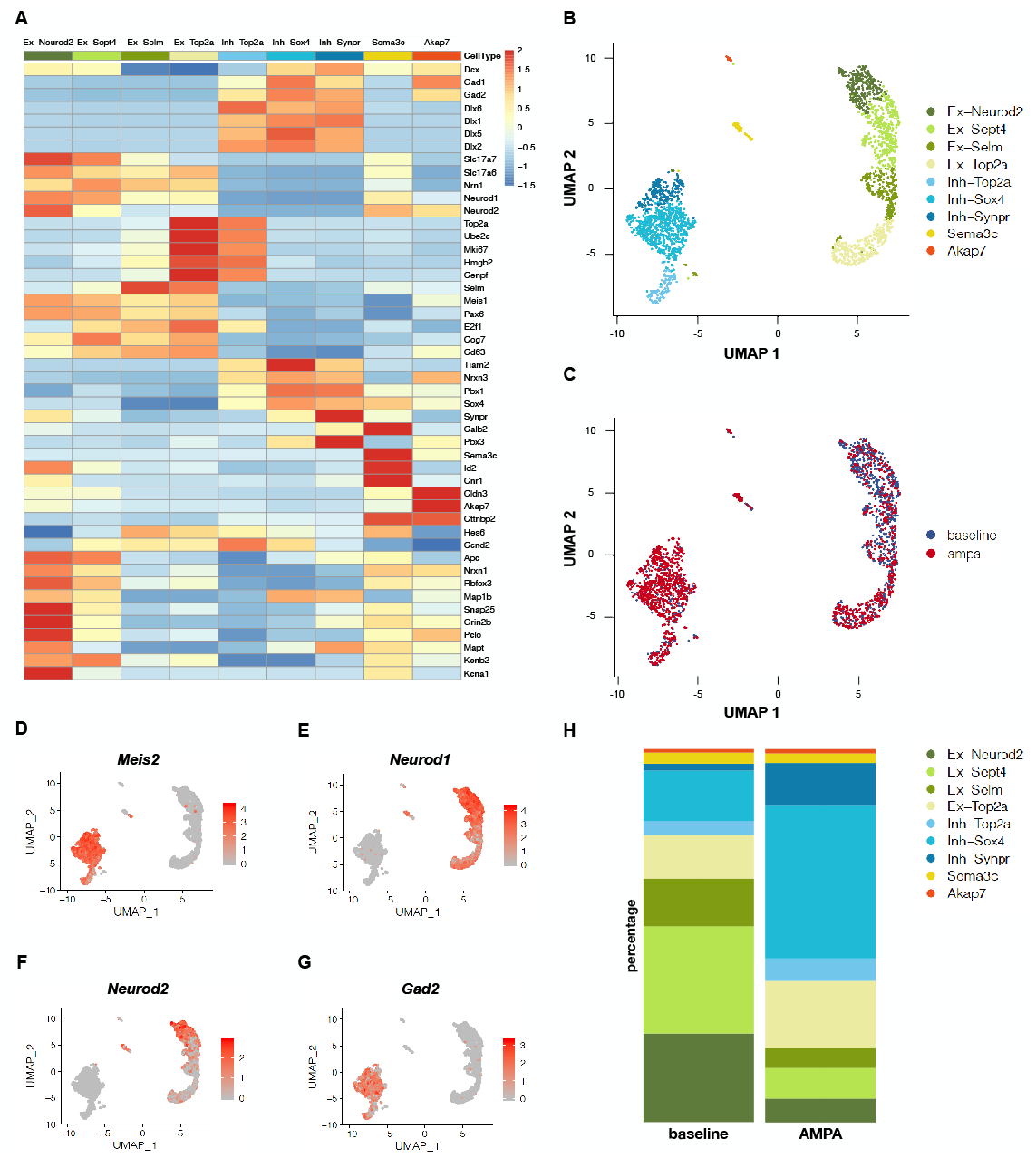
Single-cell transcriptomic analysis of GD1 cells. **A)** Heatmap of marker gene expression for the nine identified cell clusters in the aggregated dataset before and after stimulation. Ex-*Neurod2*, Ex-*Sept4*, Ex-S*elm* and Ex-*Top2a*: excitatory clusters; Inh-*Top2a*, Inh-*Sox4*, and Inh-*Synpr*: inhibitory clusters. *Sema3c*: ependymal cells. *Akap7*: oligodendrocytes. **B)** UMAP representation of the distribution of the nine clusters. **C)** UMAP representation of the two aggregated WT samples: baseline and AMPA. **D)** UMAP representation of the *Meis2*-expressing cell cluster. **E)** UMAP representation of the *Neurod1*-expressing cell cluster. **F)** UMAP representation of the *Neurod2*-expressing cell cluster. **G)** UMAP representation of the *Gad2*-expressing cell cluster. **H)** Relative proportion of cells by cluster type for individual samples.

We identified two clusters of cycling progenitors (*Top2a, Ube2c, Mki67, Hmgb2*, and *Cenpf*; Figure 3A and Supplemental Figure S2B) already engaged toward the inhibitory (*Dlx1, Dlx5* and *Dlx2*) or excitatory lineages (*Selm, Meis1, Pax6, E2f1, Cog7*, and *Cd63*). Inhibitory neurons were split in three main populations that express *Meis2* in combination with other markers: *Tiam2/Nrxn3, Pbx1/Sox4* and *Synpr/Calb2/Pbx3*. We also identified two inhibitory neuron clusters further advanced in their maturation process: “*Sema3c*” (*Calb2, Sema3c, Id2*, and *Cnr1*), composed of VIP neuron precursors, and inhibitory mature neurons composed of “*Rora*” cells (*Cldn3, Akap7, Cttnbp2, Gad1*, and *Gad2*) that may give rise to CCK interneurons (5) (Figure 3A, 3B and 3G). Next, we identified a continuum of three excitatory cell clusters, separated by their differentiation status: ventricular zone or dentate gyrus granule cell intermediate progenitors (*Hes6* and *Ccnd2*), and two clusters of further differentiated cells (*Neurod1, Neurod2, Apc, Nrxn1, Rbfox3*, and *Map1b*), that could be split according to their intermediate or final maturation, as indicated by the expression of pre- and post-synaptic markers (*Snap25, Grin2b* and *Pclo*), cytoskeletal (*Mapt*) and potassium channel expression (*Kcnb2 or Kcna1* and *Kcnk2*) (Figure 3A and 3B). Distribution of some specific markers (*Meis2, NeuroD1, NeuroD2*, and *Gad2*) is shown in the context of various clusters (Figure 3D – 3H). The post-aiFACS selection clearly shows that AMPA stimulation promoted the positive selection of *Meis2* interneurons (12 % to 38 % of the sorted cells; Figure 3H). Taking advantage of the sequencing ability to study editing, we could show that 100% of *Gria2* expressing cells carried the Q/R edited signature that happen from embryonic development onwards (not shown). This confirmes that AMPA selection with aiFACS allows to select pharmacologically competent cells, as the edited version of *Gria2* is ionotropic for sodium, the Fluo-4 detected calcium entry being consequent to cell activation depolarization.

In summary, aiFACS is a robust tool to analyze a tissue response to a pharmacological stimulation, offering new information on ion homeostasis players but also on the cellular specificities that drive the heterogeneity of the cell response.

### aiFACS selection through AMPA stimulation unveils impaired *Fmr1*-KO interneurons

In order to validate that aiFACS is helpful to study brain disorders, we applied it to dissociated neurons from mouse *Fmr1*-KO brains, the murine Fragile X Syndrome (FXS) model. This disorder is due to the lack of functioning of the Fragile X Mental Retardation Protein (FMRP). Indeed, recent studies highlighted interneuron dysfunctions in FXS (Cea-Del Rio and Huntsman 2014; Goel et al. 2018; Le Magueresse and Monyer 2013; Olmos-Serrano et al. 2010; Patel et al. 2013; Yang et al. 2018) as well as their alteration in expression, trafficking and functioning of AMPA receptors (AMPARs) (Cheng et al. 2017). To carry out this part of the study, we implemented the technique by introducing a dynamic selection to simultaneously analyze WT and *Fmr1*-KO samples. We dissociated PND 18 WT and *Fmr1*-KO brains and cells from both genotypes were stained with Fluo-4. WT cells were further labeled with Alexa-Fluor 594-coupled Wheat Germ Agglutinin (WGA 594), while *Fmr1*-KO cells were labeled with Alexa-Fluor 647 WGA. This labeling allowed the mixing of cells of both genotypes to perform a combined analysis (Figure 4A). Following the previous procedure, we selected 5,000 cells for each condition (baseline and upon AMPA stimulation).

**FIGURE 4.**
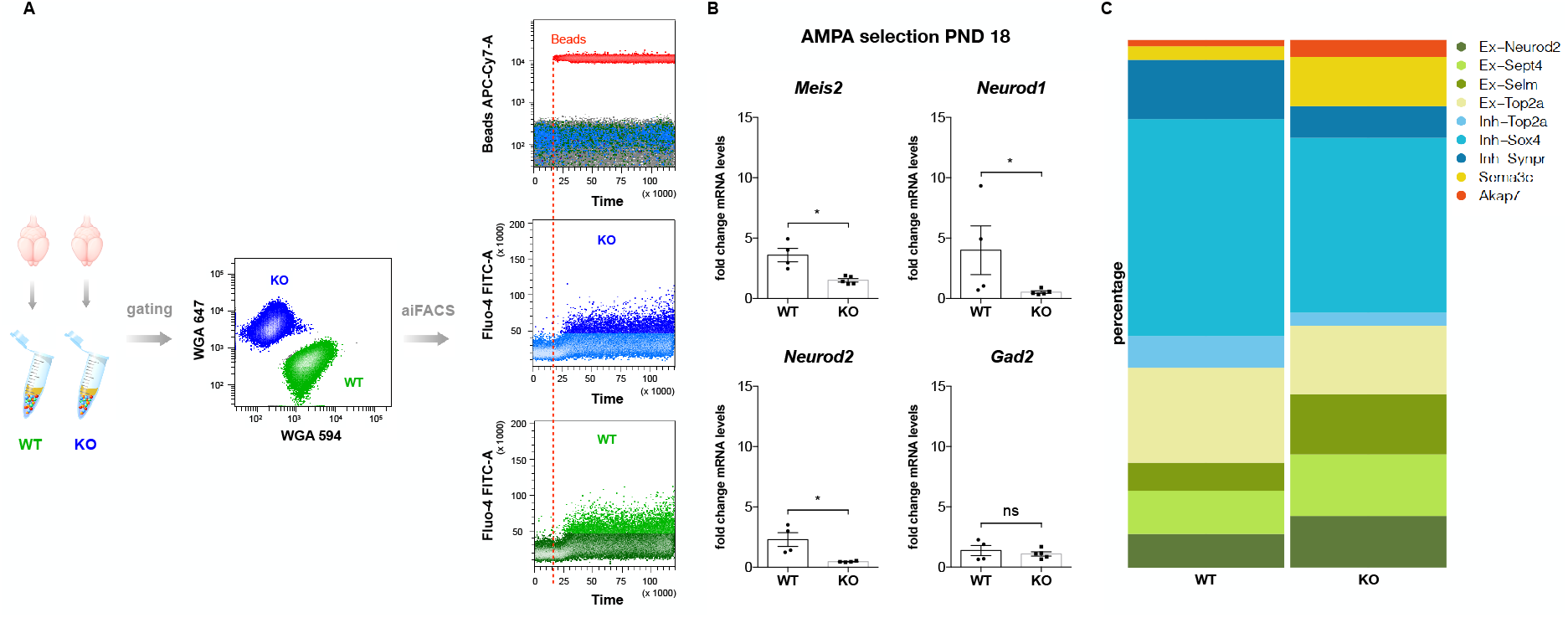
aiFACS multiplex analysis. **A)** Left panel: PND 18 WT and *Fmr1*-KO brains were processed by aiFACS. The brain dissociation is carried out mechanically with the gentleMACS™ Octo Dissociator (Miltenyi Biotec) and enzymatically with the Adult Brain Dissociation kit (Miltenyi Biotec). The selection of neuronal cells is performed magnetically with the Neuron Isolation Kit (Miltenyi Biotec). Central panel: neurons from the two genotypes were multiplexed by fluorescent labeling with Wheat Germ Agglutinin (WGA; WGA647 for WT and WGA594 for *Fmr1*-KO), processed and analyzed simultaneously. Upper right panel: the injection of fluorescently-labeled beads simultaneous to the perfusion with AMPA (100 μM final) allows to monitor the agonist concentration in the flow cell. Central and lower panels: real-time monitoring of neuronal responses to AMPA stimulation through Fluo-4 fluorescence quantification. **B)** mRNA was purified from 5,000 GD1 cells and inhibitory and excitatory marker expression levels were quantified by RT-qPCR and compared in WT and *Fmr1*-KO. Marker expression upon AMPA stimulation at PND 18 in both genotypes is presented as the fold change respective to the expression of aiFACSed WT neurons (input WT). Results are presented as the mean ± SEM, Mann-Whitney test. p values: *Meis2*, * p = 0.0159; *Neurod1* * p = 0.0317; *Neurod2* * p = 0.0286; *Gad2* ns p = 0.9048. For *Meis2*, *Neurod1* and *Gad2*: WT n = 4; *Fmr1*-KO n = 5. For *Neurod2*: WT n = 4; *Fmr1*-KO n = 4. Each n corresponds to two mouse brains and it is the mean of two independent replicates. **C)**Percentage of cells belonging to the nine clusters after single-cell analysis of AMPA response in WT and *Fmr1*-KO GD1 cells. Ex-*Neurod2*, Ex-*Sept4*, Ex-Selm and Ex-*Top2a*: excitatory clusters; Inh-*Top2a*, Inh-*Sox4*, and Inh-*Synpr*: inhibitory clusters. *Sema3c*: ependymal cells. *Akap7*: oligodendrocytes.

Since upon AMPA stimulation GD1 cells are strongly enriched in *Meis2*-expressing cells, we measured the expression levels of *Meis2* after AMPA stimulation in both WT and *Fmr1*-KO GD1 cells by evaluating the fold change respective to the expression of aiFACS selected WT neurons (Figure 4B). Our results show that the response to AMPA of GD1 *Fmr1*-KO neurons compared to WT neurons is impaired for the *Meis2*, *Neurod1* and *Neurod2* markers, while no changes were observed for *Gad2* (Figure 4B). This suggests that AMPA-responding *Fmr1*-KO GD1 cells display an abnormal level of the analyzed genes compared to WT or that a different number of cells express this marker in WT and *Fmr1*-KO. To get more insight, we performed single-cell sequencing on AMPA-stimulated GD1 cells dissociated from *Fmr1*-KO brains. We analyzed 1,047 *Fmr1-*KO GD1 cells with a median UMI per cell of 6,585 and we identified the same cell clusters present in AMPA stimulated WT GD1 cells, even if their proportions are altered (Figure 4C). This suggests an overall impairment in AMPA response in the absence of FMRP as cells from each cluster are affected. This variability could be explained by the different level of maturity or developmental profile of the cells composing each cluster in WT *vs Fmr1*-KO. To evaluate this hypothesis, we measured the levels of the analyzed markers at baseline (Figure 5A). We observed an elevated expression of the two pro-neuronal genes *Neurod1* and *Neurod2* in *Fmr1*-KO baseline GD1 cells. To assess whether the phenotype we observed is associated to the developmental process, we decided to repeat the analysis in mice older than PND 18. Remarkably, at PND 19 we observed that the expression levels of *Meis2 Neurod1*, *Neurod2* and *Gad2* do not show a significantly different abundance between the two genotypes (Mann-Whitney test, *Meis2*, ns p = 0.02857; *Neurod1* ns p = 0.2857; *Neurod2* ns p = 0.5714; *Gad2* ns p = 0.1905. Figure 5B). Consistently, the analysis of PND 19 GD1 *Fmr1*-KO neurons revealed that their response to AMPA is unchanged compared to WT neurons for all the analyzed markers (Supplemental Figure S3). We can therefore suggest that AMPA-responding *Fmr1*-KO GD1 cells might be improved at this time of development, at least for the GD1 cells expressing these markers.

**FIGURE 5.**
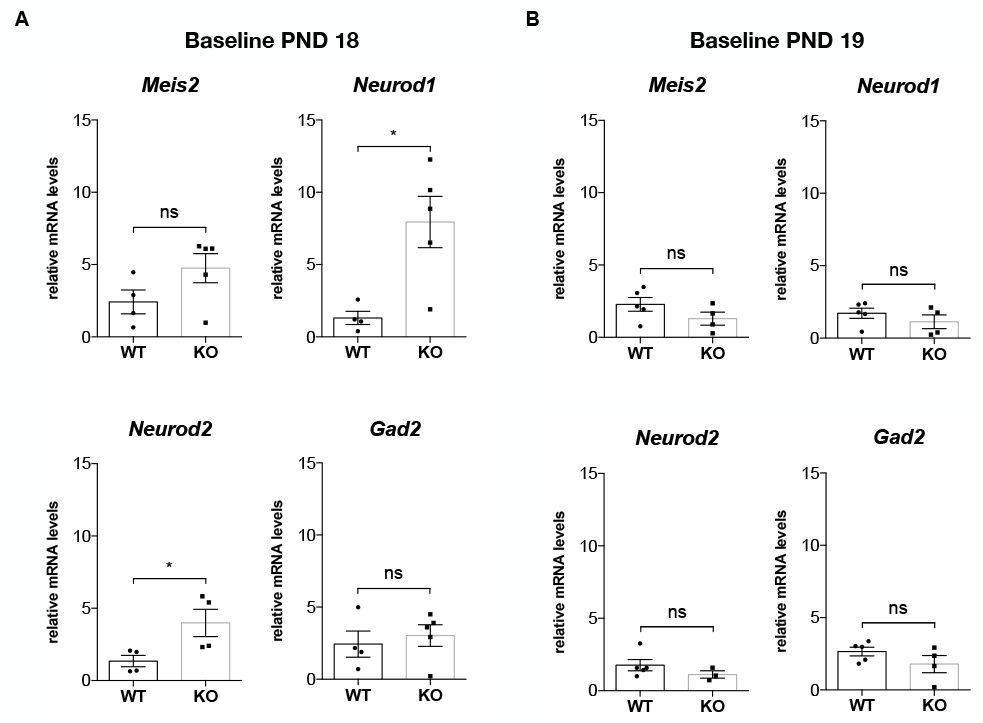
Baseline gene expression at PND 18 and PND 19. **A)** At PND18, mRNA was purified from 5,000 GD1 cells and inhibitory and excitatory marker expression levels were quantified by RT-qPCR and compared between WT and *Fmr1*-KO brains. Baseline marker expression in both genotypes is presented respective to the expression of aiFACSed WT neurons (input WT). Results are presented as the mean ± SEM, Mann-Whitney test. p values: *Meis2*, ns p = 0.1905; *Neurod1* * p = 0.00317; *Neurod2* * p = 0.0286; *Gad2* ns p = 0.7302. For *Meis2*, *Neurod1* and *Gad2*: WT n = 4; *Fmr1*-KO n = 5. For *Neurod2*: WT n = 4; *Fmr1*-KO n = 4. Each n corresponds to two mouse brains and it is the mean of two independent replicates. **B)** At PN19, mRNA was purified from 5,000 GD1 cells and inhibitory and excitatory marker expression levels were quantified by RT-qPCR and compared between WT and *Fmr1*-KO brains. Baseline marker expression in both genotypes is presented respective to the expression of aiFACSed WT neurons (input WT). Results are presented as the mean ± SEM, Mann-Whitney test. p values: *Meis2*, ns p = 0.02857; *Neurod1* ns p = 0.2857; *Neurod2* ns p = 0.5714; *Gad2* ns p = 0.1905. For *Meis2*, *Neurod1* and *Gad2*: WT n = 5; *Fmr1*-KO n = 4. For *Neurod2*: WT n = 5; *Fmr1*-KO n = 3. Each n corresponds to two mouse brains and it is the mean of two independent replicates.

## DISCUSSION

### A new approach to study pharmacological responses of cells

We developed the aiFACS prototype with the need of studying the cell-specific answer to environmental/pharmacological *stimuli*. We show here that this technique allows to sort cells according to their activity and to monitor them in real time with a fluorescent calcium indicator upon pharmacological treatment. The duration and the quality of the stimulus are the most critical aspects to manage while setting-up the aiFACS method, because they represent the key modulators of the selection. Indeed, the duration of the stimulus is dependent on the length of the red tubing (Figure 1A) and on the speed of the sample flow rate (Supplemental Figure 2A). Moreover, one or more stimuli can be applied depending on the cell line analyzed and on the cellular phenotype of interest. aiFACS-selected cells can be analyzed by various downstream methods, including single-cell sequencing. This technique allows to unravel cell identity, define the molecular determinants of the pharmacological response and directly access gene expression differences including splice variants and edited transcripts. Furthermore, by introducing dynamic selection, WT and one or more mutants can be analyzed simultaneously in the same run or multiple individuals having the same genotype can be multiplexed and analyzed separately, but simultaneously. To show the power of aiFACS, we used the brain as a highly specialized and heterogeneous tissue. By applying this technique to PND 18 mouse brains, we demonstrated the possibility of enriching our samples in interneurons. To date, the study of interneurons has been confined to restricted brain areas or circuits, due to the limited availability of high throughput analysis tools allowing a thorough and precise analysis of the complex molecular, spatial, anatomical and connection heterogeneities of the brain (Le Magueresse and Monyer 2013). By our approach, we are convinced that it will be possible to provide a novel function-based classification of interneuron cells. Of note, with more and more fluorescent reporters made available to the research community, the aiFACS can be applied to study in great detail second messenger modulations, kinase activations, ionic fluxes and many other biochemical and pharmacological mechanisms at the level of individual cells.

### The power of aiFACS to study brain disorders: the proof of concept in Fragile X syndrome

We compared WT and *Fmr1*-KO aiFACS-selected cell responses to AMPA. Our results show a global altered AMPA response in the *Fmr1*-KO cells, in agreement with other studies (Cheng et al. 2017). The same clusters of cells were selected but their ratio is different between WT and *Fmr1*-KO cells. Indeed, we validated that *Fmr1*-KO cells display abnormal levels of *Meis2*, *Neurod1* and *Neurod2*, while overall consistent levels of *Gad2* were observed (Figure 4B). These data indicate that the balance of inhibitory/excitatory neuronal activity is abnormal in the FXS brain, as expected (Contractor et al. 2015). Remarkably, we uncover the specific deregulation of *Meis2* interneuron responsiveness to AMPA stimulation. *Meis2* is a crucial transcription factor for interneuron maturation in the rodent brain. Mutations or deletions of this gene have been associated to neurodevelopmental disorders displaying ID and ASD (Giliberti et al. 2019; Verheije et al. 2019). *Meis2* is a marker of Lateral Ganglionic Eminence (LGE)-derived interneurons that are Medium Spiny Neurons (MSNs) of the striatum. Moreover, *Meis2* interneurons are in the Rostral Migratory System, giving rise to dopaminergic periglomerular interneurons of the Olfactory Bulb (OB) (Agoston et al. 2014; Allen et al. 2007; Fujiwara and Cave 2016). *Meis2* interneurons were never involved in the pathophysiology of FXS before and altered levels of *Meis2* (mRNA or protein) were never observed in studies involving large *Fmr1*-KO brain regions. Indeed, the previous analysis of cortex and hippocampus transcriptomics of *Fmr1*-KO mouse did not reveal substantial differences with WT samples (Maurin et al. 2018a). Therefore, our data clearly show that aiFACS allows to highlight unexpected impairments in cell subtypes by carrying out a functional cellular selection.

In summary, our results point out the presence of functional and molecular impairments in *Fmr1*-KO interneurons subtypes that are synchronized with brain development and they open up new research perspectives for FXS. Remarkably, we observed that the expression levels of some markers, such as the two pro-neural genes *Neurod1* and *Neurod2*, which are deregulated at PND 18, appear normalized when analyzed at PND 19. We can consider it as a consequence of an altered developmental profile of *Fmr1*-KO interneurons dependent on the function of FMRP that is known to modulate a network of pathways, which is supposed be completely disorganized in its absence (Maurin and Bardoni 2018; Maurin et al. 2018a; Miyashiro et al. 2003; Richter and Coller 2015). Remarkably, increasing evidence suggests that molecular and cellular FXS phenotypes are mitigated by age, as previously shown at the cellular or molecular levels in human and mouse brains (Nimchinsky et al. 2001; Qin et al. 2013; Tang et al. 2015; Tomasi et al. 2018). Moreover, also in ES-derived neurons we have previously shown that phenotypes present in *Fmr1* knock-down cells during the first week of *in vitro* differentiation disappeared after three weeks in culture (Khalfallah et al. 2017). Beside the molecular/cellular mechanisms underpinning those abnormalities, our data are of critical importance to define effective therapies for FXS, since we show here that PND 18 seems to be a critical development time for *Fmr1*-KO interneurons.

## Conclusions

We present aiFACS as a new tool to analyze interneurons from mouse brains. With the proof of concept on the FXS mouse model, we were able to confirm the novelty of our approach that allows the collection of a wealth of new information concerning the molecular pathology of a brain disorder. This was possible by using both a sophisticated approach such as single-cell RNA sequencing and a simple and inexpensive technique like RT-qPCR.

In perspective, it will be possible to build a novel cartography of interneurons based on their functional activities rather than on (or in addition to) other parameters such as localization, morphology and gene expression. Furthermore, by modulating the aiFACS selection parameters, testing different developmental times, using various stimuli, and multiplying the analysis of readouts, it will be possible to analyze all brain cell types both in health and in pathology. Beside the brain, future applications of aiFACS can involve the analysis of other healthy/pathological tissues, including tumors.

## METHODS

### Brain dissociation and neuron isolation

Full brains were dissected from PND 18 mice. A brief wash with complete D-PBS (supplemented with 0.5% bovine serum albumin, 1% pyruvate and 15 mM glucose) was performed before cutting the brains sagittally in six equally thick sections (2 mm) using a mouse brain matrix slicer (CellPoint Scientific) and 5 razorblades. Brain slices were dissociated mechanically using a gentleMACS™ Octo Dissociator and enzymatically using the Adult Brain Dissociation kit (Miltenyi Biotec) following the manufacturer’s instructions. The Neuron Isolation Kit (Miltenyi Biotec) was used for magnetic selection of neuronal cells.

### Neuron labelling

Neuronal suspensions were labelled at 37°C with a combination of the Fluo-4 calcium indicator (5 μg/ml; Invitrogen) and either Alexa-Fluor 594-coupled Wheat Germ Agglutinin (WGA) for WT cells or Alexa-Fluor 647-coupled WGA for *Fmr1*-KO cells (5 μg/ml; Invitrogen) in D-PBS for 20 and 10 min, respectively. A centrifugation at 300g for 10 minutes at room temperature was then performed and cells were resuspended in 300 μl of D-PBS. Prior to sorting, neurons were labeled with 0.05 μg/ml DAPI

### aiFACS and cell sorting

Cells were sorted, using a 100 μm nozzle, on a FACSAria III (BD Biosciences) equipped with four lasers. Fluo-4, Alexa Fluor 594, DAPI, Alexa Fluor 647, and APC-Cy7 (Invitrogen) were excited at 488nm, 561nm, 405nm, and 633nm respectively, and detected through BP530/30, BP610/20, BP450/40, BP660/20, and BP780/60 filters. The sorter was implemented with a homemade injection system (Figure 1A). The sample line was improved, upstream of the solenoid valve, with an injection system composed of two syringes controlled by valves and a peristaltic micro-pump. D-PBS was put in the first syringe. A 2X agonist solution (100 mM KCl or 200 μM AMPA or 10 μM ionomycin) was prepared and APC-CY7-labeled CompBead compensation particles (BD Biosciences) were added to the solution before putting it in the second injection syringe. Both the MINIPULS 3 peristaltic pump (Gilson) and the cytometer flow rate were set to 39 μl/min. Baseline acquisition and sorting were done with the valve of the buffer syringe opened. Once the valve of the agonist syringe was opened, the valve of the buffer syringe was closed. The agonist solution running in the flow cell was monitored by the appearance of the beads. In this moment the agonist-responding cells started to be sorted. Cells were collected in D-PBS. Data were analyzed with BDFACSDiva v6 software (BD Biosciences).

### RNA preparation and RT-qPCR

Total RNA from aiFACS-sorted cells was extracted using Trizol reagent (Sigma) according to the manufacturer's instructions. In each experimental sample 1μg of RNA (Supplemental Figure S1) or 5,000 cells (Fig. 2B, 4B, 5A, 5B and Supplemental Figure S3) per condition were used. RNA was purified using 500 μl of Trizol reagent and precipitated from the aqueous phase with 500 μl of Isopropanol (VWR Medicals) and 1 μl of glycogen (20 μg/μl, Invitrogen). RNA was resuspended in 20 μl (Supplemental Figure S1) or 11 μl (Fig. 2B, 4B, 5A, 5B and Supplementary Figure S3) of Nuclease Free H_2_O. Either 1 μg (Supplemental Figure S1) or 11 μl (Fig. 2B, 4B, 5A, 5B and Supplemental Figure S3) of RNA were added to the RT reaction that was performed using the Superscript IV synthesis kit (Invitrogen). An initial amplification was done with a denaturation step at 65°C for 5 min, followed by o ligo d(T) annealing at 23°C for 10 min, primer annealing at 53°C for 10 min and primer extension at 80°C for 10 min. Upon completion of the cycling steps, the reactions were stored at −20°C. Quantitative PCR (RT-qPCR) was performed on a Light Cycler 480 (Roche) with MasterMix SYBRGreen (Roche) following the manufacturer’s instructions and according to the MIQE guidelines (Bustin et al. 2009). Primer sequences are presented in Supplemental Table S1.

### Immunocytochemistry on aiFACS-sorted neurons

aiFACS-sorted neurons were plated on ornithine-coated glass coverslips (35 mm diameter) and cultivated in complete medium: Neurobasal (Invitrogen) supplemented with B-27 (Invitrogen) and GlutaMAX (Invitrogen) as previously described (Maurin et al. 2018b). Neurons were stained with the Microtubule Associated Protein 2 (MAP2; Bio Legend) antibody.

### Droplet-based scRNA-seq

Single-cell suspensions were converted to barcoded scRNA-seq libraries using the Chromium Single Cell 3’ Library, Gel Bead & Multiplex Kit and Chip Kit (10x Genomics), aiming for an estimated 2,000 cells per library and following manufacturer’s instructions. Samples were processed using kits pertaining to V2 barcoding chemistry of 10X Genomics. Libraries were sequenced on an Illumina NextSeq500, and mapped to the mouse genome (build mm10) using Cell Ranger (10X Genomics). Gene positions were annotated as per Ensembl build 84.

### Single-cell gene expression quantification and determination of the major cell types

Raw gene expression matrices generated per sample using Cell Ranger (version 2.0.0) were loaded and processed in R (version 3.4.3). Samples were analyzed independently within the Seurat workflow using the Seurat R package (version 3.0.0). First, cells that had over 95% dropouts were removed. Gene expression matrices from remaining cells were normalized using SCTransform from Seurat package. To reduce dimensionality of each dataset, the resulting variably expressed genes were summarized by principal component analysis, and the first 30 principal components further summarized using UMAP dimensionality reduction. The three samples independent analyses were then integrated using Canonical Correlation Analysis (CCA). The analysis workflow was then re-run on an integrated dataset. Cell clusters in the resulting UMAP two-dimensional representation were annotated to known biological cell types using canonical marker genes described in literature (Tasic et al. 2016; Zeisel et al. 2015, 2018).

### Statistics

The parametric t test was applied to data of two unpaired samples. Data are expressed as mean ± SEM and the p values (or adjusted p values) < 0.05 were considered as statistically significant. Statistical analysis was performed using Prism Software version 7 (GraphPad Software, Inc.).

### Animal handling and care

Animal care was conducted in accordance with the European Community Directive 2010/63/EU. WT and *Fmr1*-KO mice on a C57BL/6J congenic background were obtained from Prof. R. Willemsen (Mientjes et al. 2006) and reported as *Fmr1*-KO 2. All animals were generated and housed in groups of six in standard laboratory conditions (22°C, 55 ± 10% humidity, 12-h light/12-h dark diurnal cycles) with food and water provided *ad libitum*. Only brains from male animals were analyzed. For timed pregnancies, noon on the day of the vaginal plug was counted as E 0.5. The experiments were performed following the Animals in Research: Reporting In Vivo Experiments (ARRIVE) guidelines (Kilkenny et al. 2010). The experiments were approved by the local ethics committee (Comité d'Ethique en Expérimentation Animale CIEPAL-AZUR N. 00788.01).

## DATA AVAILABILITY

Single-cell transcriptomic data are deposited in the GEO database, accession number: GSE142274. All single-cell analyses R scripts will be made available through github (https://github.com/ucagenomix/sc.castagnola.2020).

## ACKNOWLEDGMENTS

The authors are grateful to M. Beal, M. Capovilla, M. Drozd, M. Grossi for discussion, help and encouragement. This study was supported by Inserm, CNRS, Université Côte d’Azur, Fraxa Research Foundation, Féderation Recherche sur le Cerveau, Fondation Jérôme Lejeune and Agence Nationale de la Recherche: ANR-11-LABX-0028-01, ANR-15-CE16-0015 and ANR-15-IDEX-0001. The authors acknowledge the IPMC partner “Microscopie Imagerie Côte d’Azur” (MICA) GIS-IBiSA multi-sites platform supported by the GIS IBiSA. Conseil Départemental 06, Région PACA, ARC and Canceropôle PACA and “UCAGenomiX platform », partner of the National Infrastructure France Génomique, supported by the Commissariat aux Grands Investissements: ANR-10-INBS-09-03, and ANR-10-INBS-09-02.

