## Supplementary Figures for "Agonist-induced Functional Analysis and Cell Sorting, a novel tool to select and analyze neurons: Fragile X as a proof of concept"

### SUPPLEMENTAL FIGURES

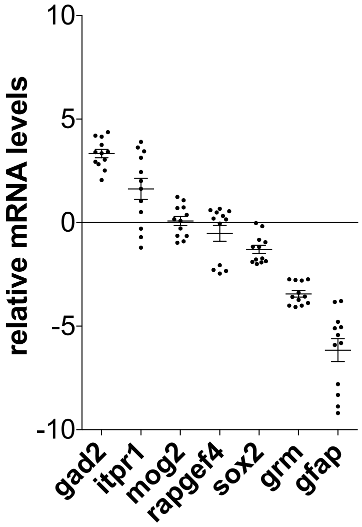


**FIGURE S1. Marker analysis of cells after dissociation.** mRNA was purified from freshly dissociated WT PND 18 mouse brains after neuronal selection. Marker levels of neuronal (*Gad2* and *Itpr1*) and non-neuronal (*Rapgef4*, *Sox2*, *Grn* and *Gfap*) cells were measured by RT-qPCR and are presented as log2 fold change respective to the non-neuronal fraction. Results are presented as the mean ± SEM. n = 12.

**A**

| Pump speed (AU) | Aria flow rate (AU) | Flow rate (µl/min) | Incubation time (sec) |
|---|---|---|---|
| 6.3 | 2 | 30 | 4.3 |
| 8.3 | 3 | 39.5 | 3.2 |
| 9.5 | 4 | 45 | 2.8 |
| 11.0 | 5 | 52 | 2.5 |
| 12.8 | 6 | 60 | 2.1 |

**B**

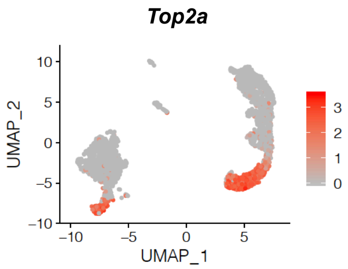


**FIGURE S2. Set-up of aiFACS with AMPA stimulation. A)** Table showing the aiFACS fluidics parameters. The speed of the peristaltic micro-pump and the flow rate of the sorter were calculated and synchronized. The parameters selected for the experiments are displayed in red. AU: arbitrary units. **B)** UMAP representation of the *Top2a*-expressing neuronal precursors, representing a less mature cell cluster in the cycling phase.

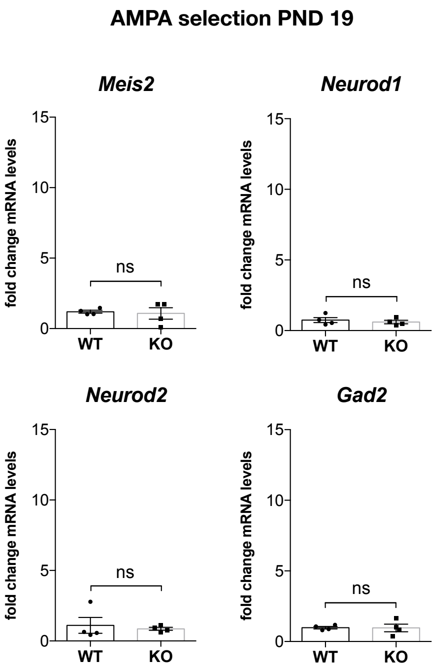


**FIGURE S3. Gene expression upon AMPA stimulation at PND 19.** mRNA was purified from 5,000 GD1 cells and inhibitory and excitatory marker expression levels were quantified by RT-qPCR and compared between WT and *Fmr1*-KO brains. Marker expression upon AMPA stimulation at PND 19 in both genotypes is presented as the fold change respective to the expression of aiFACSed WT neurons (input WT). Results are presented as the mean ± SEM, Mann-Whitney test. p values: *Meis2*, ns $p > 0.9999$; *Neurod1* ns $p = 0.6857$; *Neurod2* ns $p = 0.4857$; *Gad2* ns $p = 0.8857$. WT $n = 4$; *Fmr1*-KO $n = 4$. Each n corresponds to two mouse brains and it is the mean of two independent replicates.

| List of RT-qPCR primers (5' to 3') | | |
|---|---|---|
| **gene** | **forward** | **reverse** |
| ***Calb2*** | TGATGCTGACGGAAATGGGT | GGACATCATGCCAGAACCCT |
| ***Gad2*** | TTGATGGGAAGCCTCAACACA | AGAGGCGGCTCATTCTCTCT |
| ***Gfap*** | CAGATCCGAGGGGGCAAA | TGAGCCTGTATTGGGACAACT |
| ***Grn*** | CGTGTGCCCTGATGCTAAGA | AGATGGCATTGGGCATTGGA |
| ***Itpr1*** | GGTTCAAGCACCTGGCTACA | GGTCCACTGAGGGCTGAAAC |
| ***Meis2*** | GCCAAGGAGCAGCGTATAGT | ATTCCACTCATGGGTCCTGC |
| ***Mog2*** | GAGCTAAGAAACCCCTTTTGAGTG | GTCCACAGCAAAGAGGCCAA |
| ***Neurod1*** | ACCTTTTAACAACAGGAAGTGGA | CTCATCTGTCCAGCTTGGGG |
| ***Neurod2*** | CAAGAAGCGCGGGCCGAAGA | TTGGCCTTCTGTCGCCGCAG |
| ***Rapgef4*** | ATAAAAGGCCGTTGGAGCGA | TGACGAAGGAGGTTTGGGTG |
| ***Sox2*** | CACAACTCGGAGATCAGCA | CTCCGGGAAGCGTGTACTTA |

**TABLE S1.** List of the RT-qPCR primers used (from 5' to 3').
